# Probing individual-level structural atrophy in frontal glioma patients

**DOI:** 10.1101/2021.01.30.428923

**Authors:** Guobin Zhang, Huawei Huang, Xiaokang Zhang, Yonggang Wang, Haoyi Li, Yunyun Duan, Hongyan Chen, Yaou Liu, Bin Jing, Song Lin

## Abstract

Although every glioma patient varies in tumor size, location, histological grade and molecular biomarkers, structural abnormalities are commonly conducted in a group-level, leading to the miss of individual structural atrophy. In this study, we introduced an individual-level structural abnormality detection method for glioma patients and proposed several novel abnormality indexes to depict the individual atrophy pattern. Forty-five glioma patients in frontal lobe and fifty-two age-matched healthy controls participated in the study. All patients underwent neurocognitive test and molecular examinations, including 1p/19q co-deletion, isocitrate dehydrogenase (IDH) mutation, telomerase reverse transcriptase (TERT) promoter mutation and O6-methylguanine-DNA methyltransferase (MGMT) promotor methylation. Individual structural abnormality maps for every glioma patient were calculated from the preoperative T1 images, and the individual abnormality index were further computed and explored the associations with clinical indicators. The results manifested that: 1. Every glioma patient show unique atrophy pattern; 2. Glioma patients also share consistent atrophy regions; 3. The atrophy pattern is influenced by some molecular biomarkers. Our study provides an effective way to access the individual structural abnormalities in glioma patients, and displays great potentials in individualized precision medicine for glioma patients.

## Introduction

Gliomas represent the majority of primary central nervous system (CNS) malignant tumors, and the surgical outcomes are dependent on the comprehensive acquirement of various diagnostic information including histological grades and molecular biomarkers. However, these golden standard indicators are hard to acquire before neurosurgery, and neuroimaging has consequently served as a key way to perceive the glioma status. There are many imaging modalities collected for glioma patients, such as diffusion weighted/tensor imaging (DWI/DTI), resting-state functional MRI (rs-fMRI), positron emission tomography (PET) and structural magnetic resonance imaging (sMRI). Among them, sMRI (e.g. T1, T2) is an easy-to-collect and economical modality with high spatial resolution and test-retest reliability, which is thus widely collected in most neurosurgical centers. Usually, neurosurgeons only take qualitative evaluations for the tumor regions on structural images through visual inspection, and an individual-level quantitative analysis for tumor-related alterations is still lacking in clinical presurgical evaluation.

With the appearance of Radiomics, the quantitative analysis for the tumor region on structural images has been demonstrated to provide useful information about the histological/molecular biomarkers(Lohmann et al., 2018; Lu et al., 2018; Soike et al., 2018; Takahashi et al., 2019). However, the alterations in non-tumor regions are rarely to access in Radiomics studies, which may be also important to the postsurgical prognosis. Although the conventional morphological analysis such as voxel-based morphometry could detect structural changes (e.g. contralateral hemisphere) for glioma patients, the results are based on the group-level statistical analysis and the structural changes at individual level are still not clear. Specifically, every glioma patient may suffer unique structural alterations in non-tumor regions, and detecting and quantifying individual-level structural alterations may further deepen the understanding of the influences of tumor regions.

In recent years, individualized neuroimaging has appeared and displayed great potentials in precision diagnosis and treatment(Liu, Liu, Wang, & Dahmani, 2020; Wang et al., 2015). Stoecklein et al. found individual functional connectivity characteristics of glioma patients were significantly related with WHO tumor grade and the IDH mutation status(Stoecklein et al., 2020). However, rs-fMRI is not a clinical routine scan sequence, and to collect a high quality rs-fMRI dataset is also challenging for glioma patients. In this condition, sMRI becomes an alternative choice to model the individual alterations in the brain. Previous studies(Perry et al., 2017) have manifested W-score could successfully measure individual gray matter changes in neurological diseases, which is firstly introduced into glioma patients in the study. Furthermore, we extend the W-score method to both gray matter and white matter, and propose several novel structural abnormality indexes to depict the individual structural alteration patterns. The main purposes of the study are to ascertain: 1. Whether frontal glioma patients display unique structural alterations for every subject? 2. Whether frontal glioma patients display consensus structural alterations and where are they located? 3. Whether individual atrophy patterns vary with clinical and molecular indicators?

## Materials and Methods

### Patients

The study was approved by the Institutional Review Board of Beijing Tiantan Hospital, Capital Medical University, Beijing, China (KY2020-048-01). The study was also registered at Chinese ClinicalTrial Registry (ChiCTR2000031805). All study procedures were in accordance with the Declaration of Helsinki. Written informed consent was obtained from all patients. Fifty-two glioma patients and one hundred seventeen healthy controls participated this study, which were listed in Table 1. All gliomas in the cohort were diagnosed according to the criteria of the WHO classification system in the revised version of 2016. Inclusion criteria were suspected, newly diagnosed glioma with only one cancer region in the brain and age over 18 years. Exclusion criteria were previous cranial surgery, neuropsychiatric comorbidities, and any contraindications to MR scanning such as metal implants. Neuropsychological testing was conducted prior to the first MRI scan using the *Montreal Cognitive Assessment* (MOCA) test. Histological confirmation of the diagnosis was obtained by surgical resection. Molecular markers, including 1p/19q codeletion, IDH1/2 mutation, TERT promoter mutation and O^6^-methylguanine-DNA methyltransferase (MGMT) promotor methylation were all collected. Chromosomes 1 and 19 were analyzed by the fluorescence in situ hybridization method, and the IDH1/2 mutation and TERT promoter mutation were detected by sequence analysis, both following a previously described protocol(Suh, Kim, Jung, Choi, & Kim, 2019). MGMT promoter methylation was assessed by methylation-specific PCR as described previously by our team(Zhang et al., 2013). Patients were followed with routine clinical visits after initiation of therapy. All age-matched healthy controls were recruited from the local community and university students.

**Table 1.** The demographic and clinical information of all subjects.

|  | Glioma patients | Healthy controls |
| --- | --- | --- |
| Number | 45 | 51 |
| Gender (M/F) | 29/16 | 24/27 |
| Age | 43.2±9.7 | 42.6±9.7 |
| TIV | 1475.1±114.0 | 1425.9±144.9 |
| Tumor volume | 37.6±38.0 | - |
| Histological grade (□/□/□) | 33/10/2 | - |
| IDH1 mutation (mutated/wild) | 39/6 | - |
| TERT promoter mutation<br>(mutated/wild) | 33/19 | - |
| 1p/19q deletion (both 1p/19q /only<br>19q/only 1p/none) | 24/2/4/15 | - |
| MGMT promoter methylation | 0.25±0.16 | - |
| MOCA | 22.6±4.6 | - |

Abbreviation: IDH, isocitrate dehydrogenase; MGMT, O<sup>6</sup>-methylguanine-DNA methyltransferase; MOCA, Montreal Cognitive Assessment; TERT, telomerase reverse transcriptase; TIV, total intracranial volume.

### Structural MRI acquirement

All subjects were scanned with a Philips Ingenia 3.0T MRI scanner at Beijing Tiantan Hospital. For both glioma patients and healthy controls, T1 sequence was collected with the following parameters: TR: 6.5 ms, TE: 3.0 ms, Flip angle: 8°, voxel size: 1×1×1 mm^3^, image dimension: 256×256×196. In addition, T2-flair was also scanned for glioma patients with the listed parameters: TR: 4.8 s, TE: 0.34 s, Flip angle: 90°, voxel size: 0.625×0.625×0.55 mm^3^, image dimension: 400×400×300.

### Image processing

The tumor region of every patient was extracted from T2-flair images with two sequential steps: 1. launch an automatic segmentation by ITK-SNAP software; 2. manually correct the segmentation by experienced neurosurgeons. After the segmentation, individual T2-flair image was co-registered to its corresponding T1 image by SPM software, and the transformation matrix was used on the segmented tumor to obtain the matched tumor region in T1 image space.

Individual T1 images were processed with CAT12 software to calculate the brain tissue volume. The skull stripping and correction for bias-field inhomogeneities were subsequently conducted. After that, the whole brain was segmented into different tissue types, e.g. gray matter and white matter. Then, the segmented GM and WM images were normalized to the MNI standard space with a modulation manner by DARTEL algorithm. Finally, the normalized GM and WM images were smoothed with 4-mm FWHM Gaussian kernel.

### Individual structural atrophy map

Figure 1 illustrated the flowchart to generate individual structural atrophy map. First of all, all healthy controls were used to construct the normative brain volume model. In this model, general linear model (GLM) was adopted to discover the relationship between voxel-wise volume and variables including age, sex, and total intracranial volume (TIV) as the following equation.

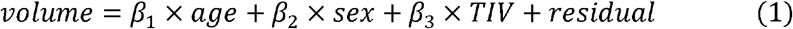

**Figure 1.**
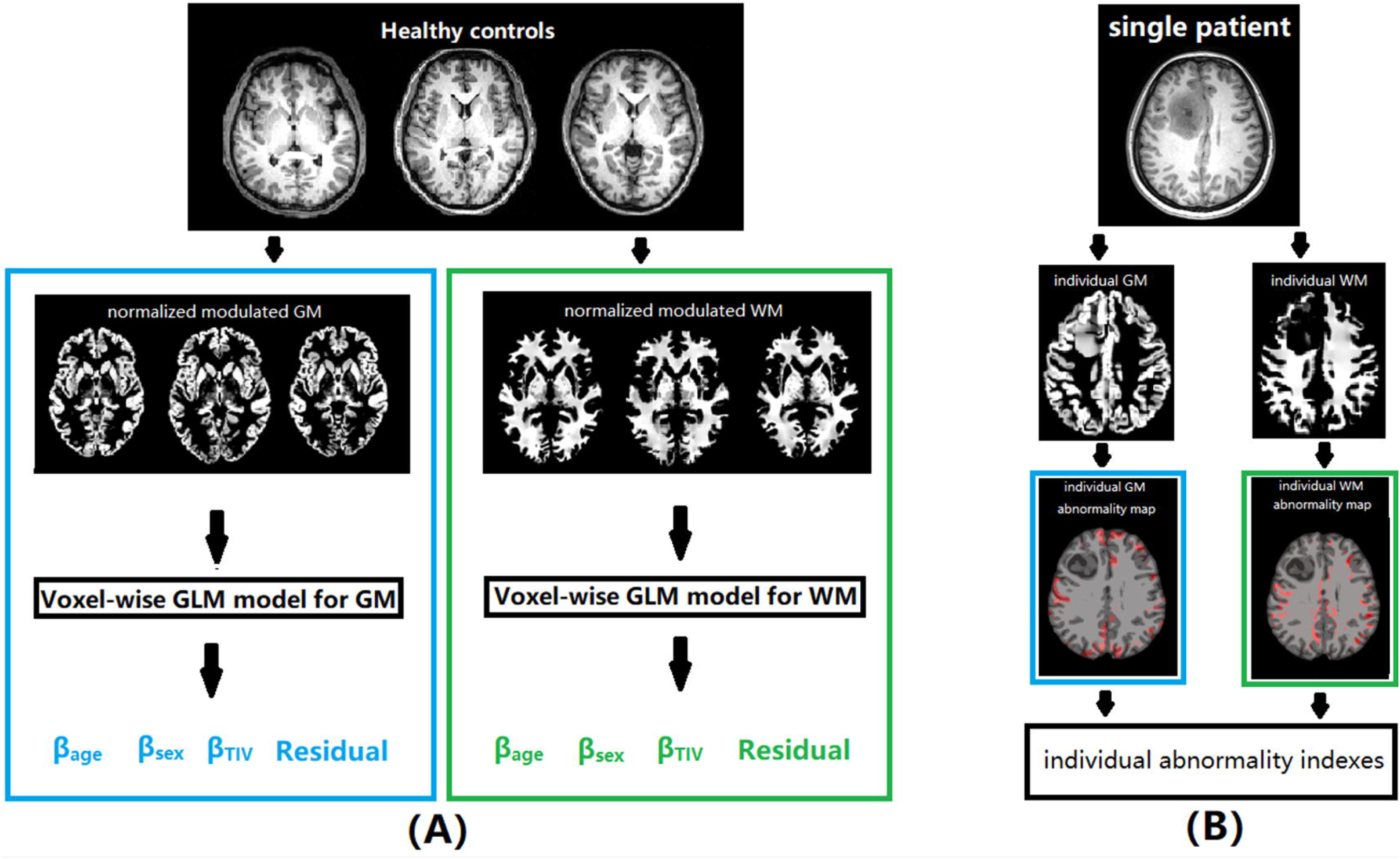
The flowchart of the proposed structural abnormality indexes: (A). the construction of normative brain volume model; (B). the calculation of the proposed individual abnormality indexes.

Here, β_1_, β_2_, β_3_ were weights for age, sex and TIV on the voxel-wise volume, and GM and WM volume models were respectively constructed.

Once the models had been created, the individual structural abnormality map for each glioma patient was calculated based on W score, which was calculated as following:

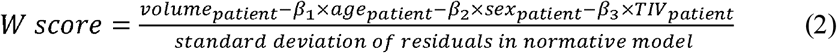

After the calculation of W score for each patient, a cutoff threshold (|W|>6) and a cluster size (K>100) were set to generate the individual structural abnormality map. Notably, an individualized explicit non-tumor mask was produced for every patient in which individual tumor mask was deleted from the tissue (GM/WM) prior probability template (threshold: 0.2). Based on the individual structural abnormality map, we further proposed 4 novel abnormality indexes on GM/WM to reflect the characteristics of structural damages induced by glioma: Ipsilateral atrophy ratio, Contralateral atrophy ratio, Non-cancer atrophy ratio, Relative atrophy ratio, which were listed in details in Table 2.

**Table 2.** The proposed 8 structural abnormality indexes.

| Abnormality index | Description |
| --- | --- |
| Ipsilateral atrophy ratio | $= \frac{\text{number of GM/WM atrophy voxels in hemisphere with tumor}}{\text{number of all voxels in hemisphere with tumor}}$ |
| Contralateral atrophy ratio | $= \frac{\text{number of GM/WM atrophy voxels in contralateral hemisphere}}{\text{number of all voxels in contralateral hemisphere}}$ |
| Non-cancer atrophy ratio | $= \frac{\text{number of abnormal GM/WM voxels in non – cancer region}}{\text{number of all voxels in whole brain}}$ |
| Relative atrophy ratio | $= \frac{\text{number of abnormal GM/WM voxels in non – cancer region}}{\text{number of all voxels in cancer region}}$ |

Abbreviation: GM, gray matter; WM, white matter.

### Associations with various glioma indicators

To explore the associations between each of 8 abnormality indexes and various glioma indicators, for continuous variables like tumor volume, MGMT methylation, and MOCA score, Spearman correlation was used to ascertain the linear relationship with these atrophy indexes; for categorical variables such as TERT mutation, 1p/19q deletion, histological grade and IDH1, two-sample T test was used to determine whether the atrophy indexes varied with these indicators. An FDR corrected P <0.05 was thought as the significance level for both analyses.

## Results

### The clinical characteristics for glioma patients

There were 45 glioma patients (mean age 43.2±9.7 years, 29 male) and 51 healthy controls (mean age 42.6±9.7 years, 24 male) that participated in the study. All patients were prospectively included between May 2019 and July 2020. The WHO histological grade, IDH1, TERT mutation, 1p/19q deletion, MGMT methylation were acquired for every patient, and Montreal Cognitive Assessment (MOCA) score was also recorded. Table 1 summarized the demographic and clinical information for all subjects.

### Every glioma patient displayed distinct atrophy pattern

All patients only displayed atrophies in GM and WM. Figure 2 illustrated three selected patients’ T1 images and corresponding individualized GM/WM atrophy maps (mapping back into T1 space). Obviously, the structural atrophies not only lay in regions close to tumors but also in regions far away from tumors. Although the patients vary in tumor sizes and tumor grades, it is clear that every subject displayed unique atrophy pattern.

**Figure 2.**
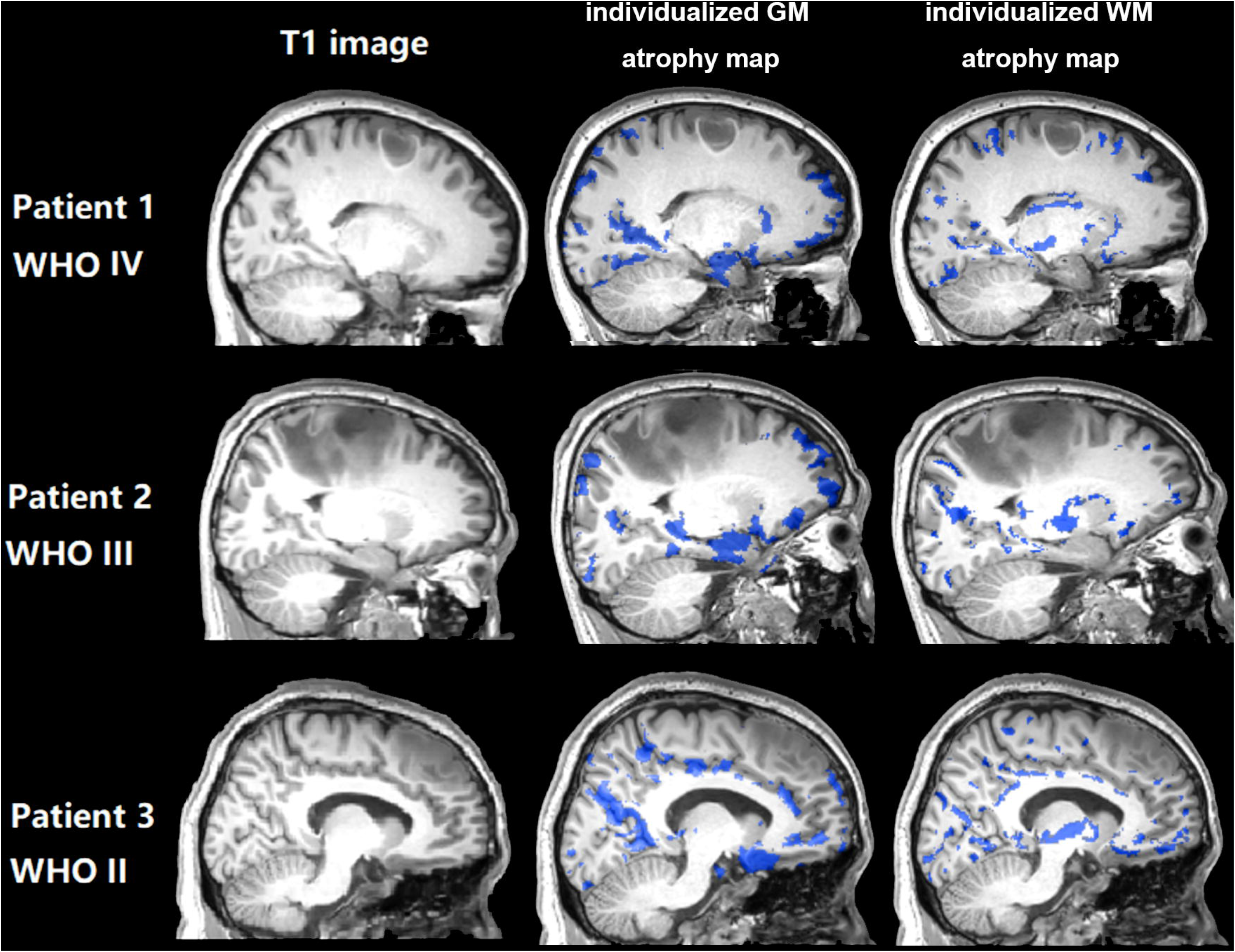
Three examples of individualized GM/WM abnormality maps.

### Every glioma patient also showed common atrophy regions

Figure 3 displayed the overlapping GM regions in individual structural atrophy map. No matter the tumor is located at the left hemisphere or right hemisphere, the atrophies in right temporal lobe are more obvious than the left one, mainly including hippocampus, amygdala, parahippocampus and thalamus. Moreover, the contralateral frontal regions also displayed some degree of atrophy. Figure 4 exemplified the consistent WM regions in individual structural atrophy map. The main atrophy regions are located at bilateral thalamus and pallidum, and the contralateral part is more severely atrophied than the ipsilateral part.

**Figure 3.**
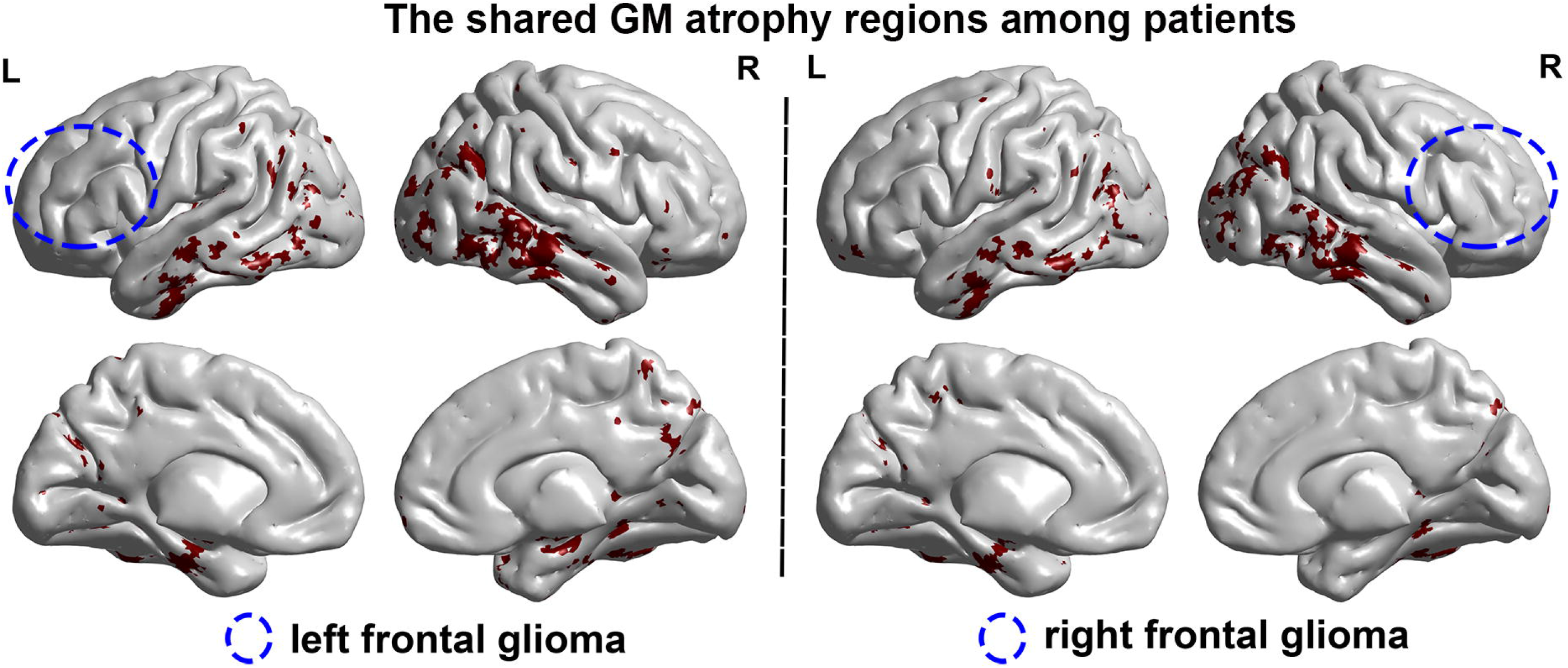
Overlapping gray matter atrophy in non-tumor regions for all patients with tumor on (A) left hemisphere and (B) right hemisphere.

**Figure 4.**
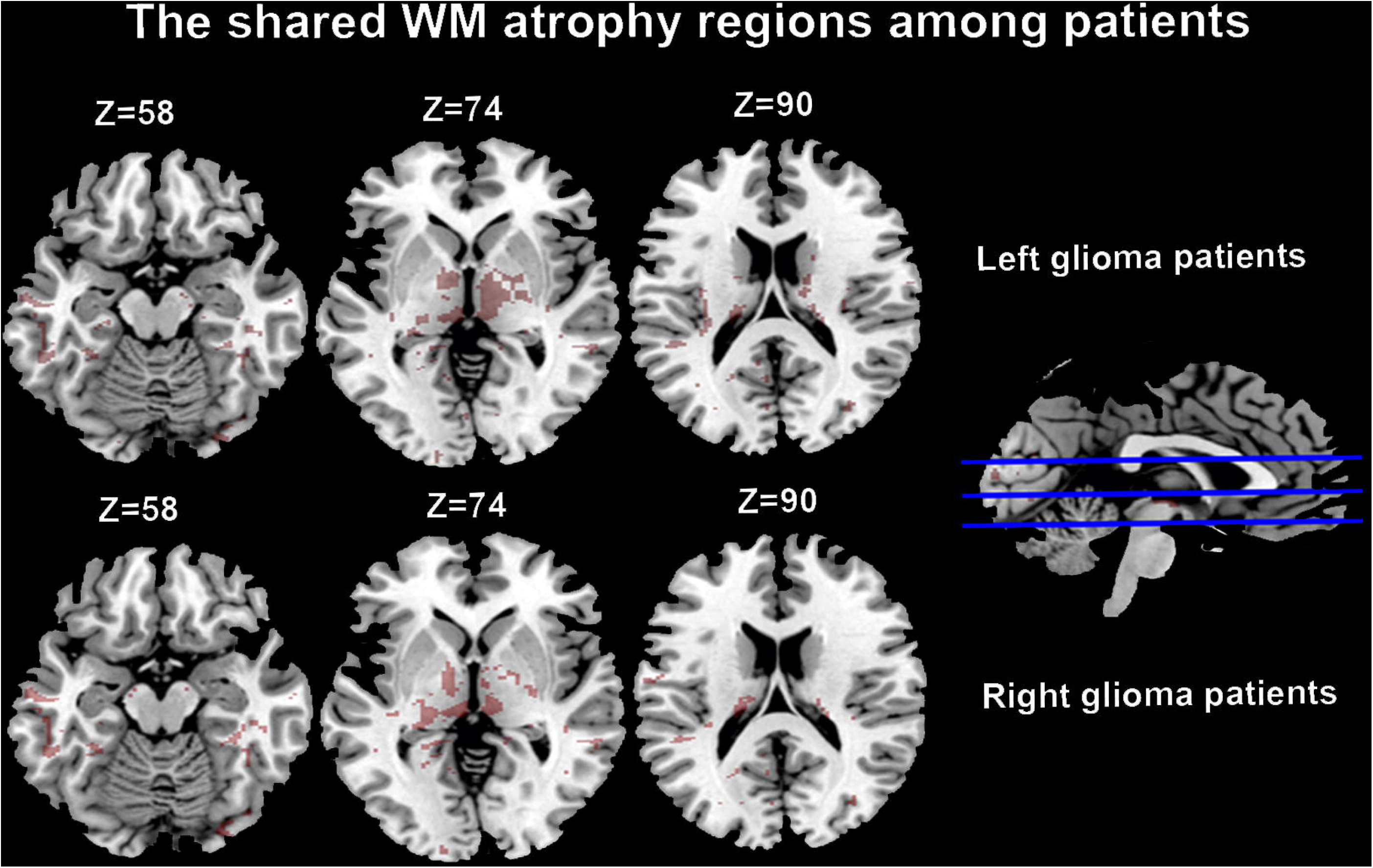
Overlapping white matter atrophy in non-tumor regions for all patients with tumor on left hemisphere and right hemisphere.

### Individual brain atrophy indexes were associated with some clinical indicators

Fig 5 illustrated all possible relations between the proposed individual abnormality indexes and clinical indicators, and all of them had passed an FDR correction with P<0.05. GM/WM relative atrophy ratio were found to significantly correlate with the tumor size, however, with the increasement of the tumor volume, the relative atrophy ratio was decreased (power law distribution), and no other individual structural indexes were related with tumor size. WM contralateral atrophy ratio was found with significant differences between IDH wild type and mutation type, while WM relative atrophy ratio were obviously different between 1p/19q wild type and mutation type. Moreover, both GM/WM relative atrophy ratio displayed between-group differences in TERT wild type and mutation type. No other relationships were detected with MOCA and MGMT.

**Figure 5.**
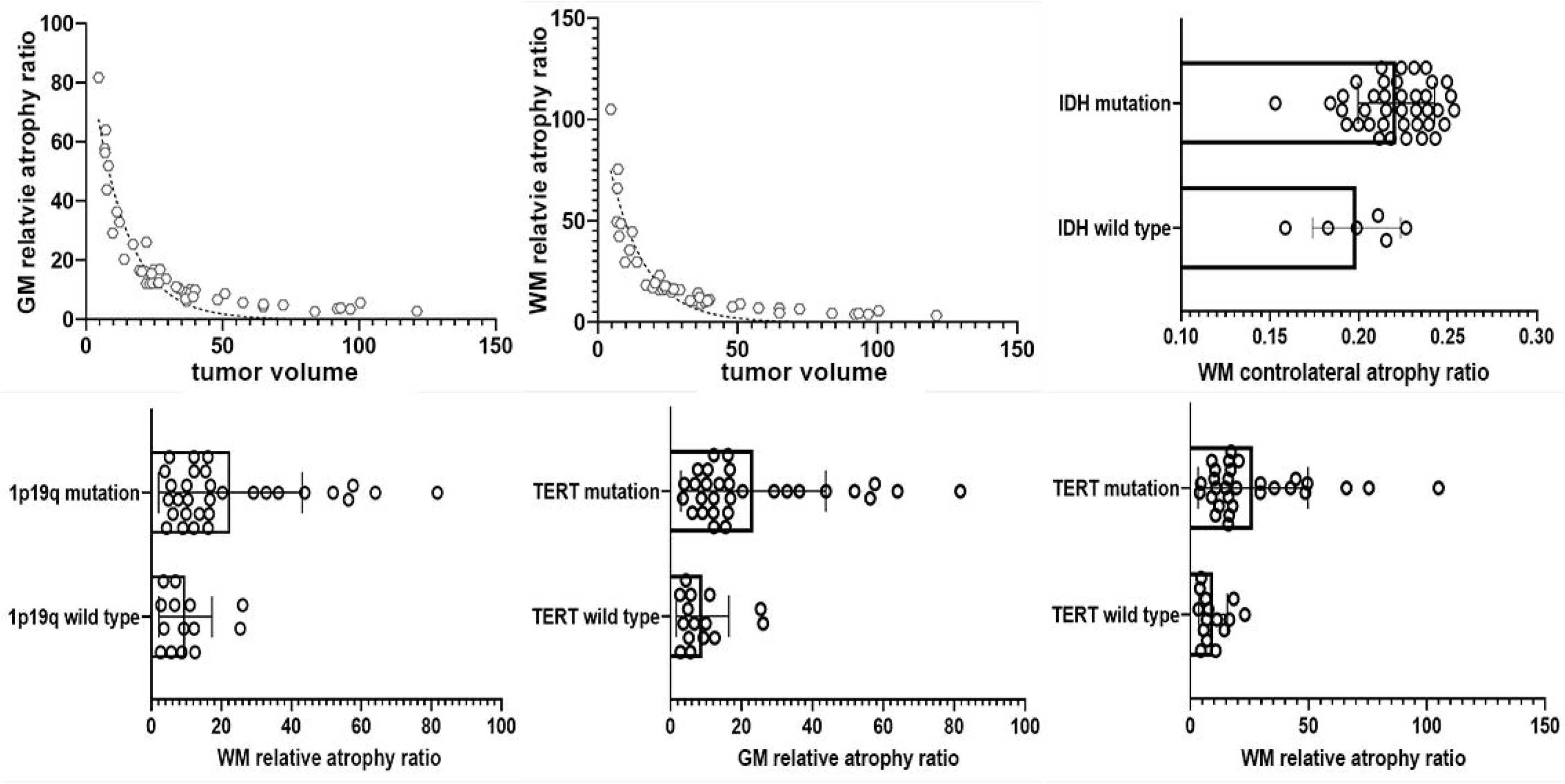
The relationship between abnormality indexes and clinical/molecular indicators.

## Discussion

In this study, we firstly depicted individual non-tumor structural atrophy characteristics for glioma patients, and proposed several quantitative indexes to ascertain the relationship between individual atrophy patterns and clinical indicators. The results demonstrated every glioma patient displays distinct atrophy pattern but also shared overlapping atrophy regions. In addition, several atrophy indexes were related with tumor size and molecular indicators.

Neuroimaging has acted as indispensable tool for presurgical assessment of glioma patients. However, most imaging studies only focus on the tumor region in order to judge the status of tumor, and the non-tumor alterations are rarely to access. Moreover, neuroimaging is usually used by two conventional manner: 1. visual inspection by experienced neurosurgeons; 2. group-level statistical comparison between glioma patients and healthy controls. Both manners may miss the individual unique changes in non-tumor regions. Our study firstly offers an individual-level manner to describe individual atrophy in every glioma patient, and manifested that every glioma patient displays unique structural atrophy in non-tumor regions, which was consistent with previous individual fMRI study(Stoecklein et al., 2020). Taken together, glioma may lead to patient-specific alterations in both brain function and structure, and individualized neuroimaging methods show great potentials in the pre-surgical evaluation and post-surgical prognosis estimation.

Although glioma patients display distinct structural atrophy at the individual level, it is interesting to find they also share overlapping GM atrophies in regions like bilateral hippocampus, amygdala, thalamus and parahippocampus. Specially, these atrophies are not dependent on the hemisphere of tumor, tumor size, histological grade or molecular indicators, indicating that frontal glioma could introduce systematic atrophy in brain structure. In addition, GM atrophy were more obvious in right temporal lobe than the left one regardless the tumor hemisphere, implying right temporal lobe is more vulnerable to the frontal tumor, and there is lateralization in the structural damages induced by the frontal tumor. Previous literatures have reported that as many as 90% of brain tumor patients would show tumor-related cognitive deficits (e.g., memory, attention, information processing, executive function)(Gehring, Roukema, & Sitskoorn, 2012; Gehring, Sitskoorn, Aaronson, & Taphoorn, 2008), which may be explained by the current findings that limbic system including hippocampus, amygdala, parahippocampus and thalamus are damaged in glioma patients.

The consistent WM atrophies were mainly located at bilateral thalamus and pallidum, and the contralateral hemisphere was more severely atrophied than the ipsilateral one. These regions are involved in two fiber bundles linking fronto-temporal regions: superior longitudinal fasciculus (SLF) and uncinate fasciculus (UF). SLF has been reported to correlate with the spatial working memory deficit(Kinoshita et al., 2016), visuospatial dysfunction(Nakajima et al., 2017) and transmission of speech and language(Henderson, Abdullah, Verma, & Brem, 2020) in glioma patients. UF has been demonstrated to relate with the language(Duffau, Gatignol, Moritz-Gasser, & Mandonnet, 2009) and cognitive deficits(Incekara, Satoer, Visch-Brink, Vincent, & Smits, 2018) in glioma patients. The common WM atrophies also prompt the possible cognitive damages in frontal glioma patients, and may provide a possible pathway that link the remote GM atrophy covariation.

GM/WM relative atrophy ratio but not other individual indexes were negatively related (in fact power law distribution) with tumor volume, and two conclusions may thereby be generated: 1. the volume of atrophy in the brain was in fact not dependent on the tumor volume; 2. the relative atrophy volume becomes smaller when the tumor volume enlarges. Moreover, WM contralateral/relative atrophy ratio were found with significant differences in IDH and 1p/19q mutation. IDH and 1p/19q are both known indicators for the prognosis of glioma patients. For example, lower grade gliomas with wild-type IDH were reported to show similar prognosis with glioblastomas, and IDH mutated glioblastomas were found with better prognosis than IDH wild-type glioblastomas(Darvishi et al., 2020; Hartmann et al., 2010). In addition, anaplastic gliomas with IDH wild-type have worse prognosis than glioblastomas with IDH mutation(Hartmann et al., 2010). 1p/19q co-deletion is found with better prognosis for patients with oligodendroglioma after radiotherapy or alkylating chemotherapy(Jenkins et al., 2006; van der Voort et al., 2019). Additionally, GM/WM relative atrophy ratio were found differences in TERT mutation, which is also a promising indicator for the treatment response of radiotherapy and temozolomide in primary glioblastoma multiforme (GBM)(Eckel-Passow et al., 2015; Peng et al., 2020; Vuong et al., 2017; Yang et al., 2016; Yuan et al., 2016). In summary, individual structural atrophy patterns are influenced by the genetic type.

Finally, several limitations should be mentioned: 1. The optimal threshold for W is not clear, and we chose |W|>6 in order to reduce the false positive rate in atrophy detection. 2. The gender and TIV are not well matched between glioma patients and healthy controls, and future studies could involve more patients from multicenter to verify the robustness of the current findings.

## Conclusion

Our study firstly depicted the individual atrophy pattern for glioma patients, and found individual structural atrophy was unique and driven by genetic information. Our findings could provide valuable information for the individualized presurgical evaluation and postsurgical prognosis.

## Data Availability

The data that support the findings of this study are available from the corresponding author upon reasonable request.

## Funding

This work was supported by the National Natural Science Foundation of China (81801042), Beijing Hospital Authority Youth Programme (20190504), Beijing Outstanding Talent Training Foundation, Youth Backbone Individual Project (2018000021469G230) and Beijing Municipal Commission of Education (KM202010025025). The funds had no role in the study design, data collection and analysis, decision to publish, or manuscript preparation.

## Conflict of Interest

The authors declare that they have no competing interests.

## Ethics approval

The study was approved by the Institutional Review Board of Beijing Tiantan Hospital, Capital Medical University, Beijing, China (KY2020-048-01).

## Patient consent

Written informed consent was obtained from all patients.

## Acknowledgments

None.

## Notes

### Competing Interest Statement

The authors have declared no competing interest.

### Summary of Updates

We update the method to investigate brain alteration in frontal lobe glioma patients, and found some new results.

